# High-throughput method to test antimicrobial gels against a multispecies oral biofilm

**DOI:** 10.1101/2020.09.08.287391

**Authors:** Kanchana Chathoth, Bénédicte Martin, Martine Bonnaure-Mallet, Christine Baysse

## Abstract

Periodontitis, characterized by the damage of the periodontium can eventually lead to tooth loss. Moreover, severe forms of periodontitis are associated with several systemic disorders. The evolution of the disease is linked to the pathogenic switch of the oral microbiota comprising of commensal colonizers and anaerobic pathogens. Treatment with antimicrobial gels has the potential to help eradicate periodontal pathogens. Testing antibacterial gels against *in vitro* biofilm models is complicated. Recovery of detached and sessile bacteria from *in vitro* biofilms treated with gel formulations using conventional methods (microtiter plates, μ-slides, flow cells etc.,) may prove arduous. To overcome this challenge, we optimised a simple method using the principle of the Calgary Biofilm Device (CBD) for testing antimicrobial gels against multispecies oral biofilms. First, we established three-species oral biofilms consisting of two periodontal pathogens (*Porphyromonas gingivalis*, *Treponema denticola*) and a primary colonizer of the dental plaque (*Streptococcus gordonii*) on the surface of pegs. Next, a protocol to test gels against oral biofilms was implemented using commercially available gels with different active products. This method enables the analysis of the composition of biofilm and detached/planktonic cells to measure the effect of topical gel formulations/antibacterial gels for the treatment of periodontitis. However, the method is not restricted to oral biofilms and can be adapted for other biofilm-related studies.

## INTRODUCTION

Periodontitis is a polymicrobial chronic inflammatory disease of the periodontium caused by the accumulation of dental plaque (Pihlstrom *et al.*, 2005). It is regarded as the second most common disease worldwide, with severe forms affecting up to 10 to 15% of adults (Petersen and Ogawa, 2012) resulting in tooth loss. Many studies have reported an association of severe periodontitis with several systemic disorders (Beukers *et al.*, 2017; Bui *et al.*, 2019; Graves *et al.*, 2019; Hujoel *et al.*, 2003; Koziel *et al.*, 2014; López, 2008; Preshaw *et al.*, 2012) and oral malodor (Lee *et al.*, 2003; Yaegaki, 2008). Therefore, periodontal diseases are a major socio-economic concern.

Metagenomics analysis of the microbial community in the human subgingival plaque has revealed the presence of over 500 species (Ai *et al.*, 2017). This oral biofilm formation is initiated by early colonizers that recognize receptors in the acquired pellicle that coats the enamel of the tooth. These early colonizers consist mainly of facultative anaerobic Gram-positive bacteria such as *Streptococcus* spp. and *Actinomyces* spp. In the oral health-associated biofilm, these Gram-positive cocci and rods predominate. Change in dominant species with an increase of putative pathogens like *Porphyromonas gingivalis*, *Treponema denticola*, *Tannerella forsythia* and *Prevotella intermedia* results in dysbiosis and thus periodontal disease (Kolenbrander *et al.*, 2002; Meuric *et al.*, 2017). The mechanism behind the shift from health-associated oral microbiota to periodontal pathogens is not clearly understood. This pathogenic shift is probably linked to the changes in the composition and/or virulence of microbiota as a result of changes in the oral environment (Pöllänen *et al.*, 2013). In our recent review (Chathoth *et al.*, 2020), we hypothesize that an excess of iron and the resultant ROS generated in presence of the peroxygenic streptococci may be one of the contributors for dysbiosis. Therefore, it is of interest to be able to access changes in biofilm composition in response to a treatment.

Antimicrobial agents in the form of gel formulations are a promising delivery system for the treatment of periodontitis *via* topical administration. The advantages include the ease of use, increased retention time at the site of application and controlled drug release. Several authors have demonstrated the effectiveness of gel formulations in reducing microbial content or plaque index (Figueiredo de Almeida Gomes *et al.*, 2006; Noyan *et al.*, 1997; Paquette *et al.*, 1997; Sauvêtre *et al.*, 1993) in human, animal-based or *in vitro* studies. Similar improvement in probing depth and/or bleeding was reported (Esposito *et al.*, 1996; Graça *et al.*, 1997) on use of gel formulations alone or in conjunction with other modes of treatment. There is an increasing interest in the use of biodegradable and biocompatible compounds like chitosan (İkinci *et al.*, 2002), poly(lactic-co-glycolic acid)/hyaluronic acid (Noda *et al.*, 2018), cranberry juice concentrate (H.R. *et al.*, 2017) in gel formulations. The non-toxic properties of such compounds combined with the advantages of gel as a delivery system is continuously explored as a treatment strategy.

Conventional methods for growing biofilms *in vitro* (microtiter plates, μ-slides, flow cells etc.) pose several difficulties in testing antimicrobial gels against biofilms. The biofilm and the treatment ought to be in the same place. Owing to the viscosity and adhesive nature of gels, the recovery of detached and sessile bacteria for assessment after treatment gets complicated. The Minimum Biofilm Elimination Concentration (MBEC™) Assay System (formerly the Calgary Biofilm Device) was previously challenged with gel-based products to assess their bactericidal activity on mono-species biofilm (Martineau and Dosch, 2007; Santos *et al.*, 2016). Hence, a method adapted to both gel-based products and polymicrobial biofilms, and capable of deciphering the behaviour of each species in response to the treatment was needed. In this study, we combined a method using the principle of the Calgary Biofilm Device (CBD) (Ceri *et al.*, 1999) and a new medium for oral bacteria (Martin *et al.*, 2018) for testing antimicrobial gels against multispecies oral biofilms. This protocol using a lid with pegs and a 96-well microtiter plate was adapted to establish three-species oral biofilms on the surface of the pegs. The three species consisted of two key periodontal pathogens *P. gingivalis*, *T. denticola* and a primary colonizer of the dental plaque, *S. gordonii*. The basis for the selection of these microorganisms was the species-specific co-aggregation of *S. gordonii* with *P. gingivalis* (Lamont and Hajishengallis, 2015) and the syntrophy and synergy between *P. gingivalis* and *T. denticola* (Meuric *et al.*, 2013; Tan *et al.*, 2014; Zhu *et al.*, 2013). Moreover, *P. gingivalis* and *T. denticola,* co-exist in deep periodontal pockets (Kigure *et al.*, 1995; Kumawat *et al.*, 2016) and are associated with severe forms of periodontitis. *S. gordonii* is a peroxygenic bacteria and a glutathione producer that can also influence the pathogenic switch of the oral subgingival biofilm (Chathoth *et al.*, 2020). Most importantly, we use saliva-coated pegs to grow the three-species biofilm *in vitro* to mimic the dental plaque developing initially at the root of teeth. This biofilm was realised in the MMBC-3 medium which allows the growth of the three bacterial species (Martin *et al.*, 2018). The method was adapted to analyse the composition of the biofilm and detached planktonic growth in order to test topical gel formulations/antibacterial gels against oral biofilms for the treatment of periodontitis.

The method was challenged with two commercial gels, Hyalugel^®^-ADO (Ricerfarma, Milan, Italy) and blue^®^m oral gel (blue^®^m Europe B.V., Netherlands). Hyalugel^®^-ADO mainly consists of hyaluronic acid (0.2 %) which is a major component of the extracellular matrix of the skin and plays a vital role in skin repair (Neuman *et al.*, 2015). Hyaluronic acid showed an antibacterial activity against both planktonic bacteria and biofilms (Ardizzoni *et al.*, 2011; Binshabaib *et al.*, 2020; Eick *et al.*, 2013; Pirnazar *et al.*, 1999). The main ingredients of the blue^®^m oral gel are sodium perborate (1.72 %) and lactoferrin (0.2 %). Sodium perborate acts as the oxygen donor that can be lethal to the anaerobic periodontal pathogens. Lactoferrin has antimicrobial, anti-inflammatory and anti-carcinogenic properties and also acts as an iron chelator (Wang *et al.*, 2019). Previous reports have demonstrated that blue^®^m oral gel reduced *P. gingivalis* planktonic growth (Deliberador *et al.*, 2020) and also showed antiplaque and anti-gingivitis efficacy in the form of a toothpaste (Cunha *et al.*, 2019).

## METHODS

### Strains and media

Strains of *Streptococcus gordonii* Challis DL1 (Chen *et al.*, 2004), *Porphyromonas gingivalis* TDC60 (Watanabe *et al.*, 2011) and *Treponema denticola* ATCC35405 (Chan *et al.*, 1993) were used for the study. The cultures of *S. gordonii* and *P. gingivalis* were grown in MMBC-3 (Medium for Mixed Bacterial Community), with FeSO_4_.7H_2_O_2_ (Sigma-Aldrich) at 8 μM and protoporphyrin IX (PPIX, Sigma-Aldrich) at 0.08 μM as the iron source (Martin *et al.*, 2018). *T. denticola* was initially grown in NOS spirochete medium (Leschine and Canale-Parola, 1980) and further sub-cultured in MMBC-3 supplemented with FeSO_4_.7H_2_O_2_ (8 μM) and PPIX (0.08 μM). All three micro-organisms were grown in anaerobic condition at 37°C in an anaerobic chamber (MACS 500, Don Whitley Scientific, United Kingdom) with 10% v:v H_2_, 10% v:v CO_2_ and 80%v:v N_2_.

### Growth and treatment of the biofilm

The protocol for the growth and treatment of the biofilm is detailed in Figure 1. As the first step, 200 μl of saliva (Pool Human Donors, MyBioSource), filtered (0.20 μm) and diluted twice in sterile distilled water, was added into the Nunc™ Nunclon™ 96-well tissue culture microtiter plates. The lid with pegs (Nunc-TSP, polystyrene) was placed over the microtiter plate and incubated in saliva for 30 minutes. Next, the saliva was replaced by 200 μl of inoculum consisting of *S. gordonii* (OD_600nm_:0.05), *P. gingivalis* (OD_600nm_:0.1) and *T. denticola* (OD_600nm_:0.1) in a new microtiter plate. The lid with pegs was placed on the microtiter plate ensuring that the pegs were immersed in the inoculum. This set-up was incubated in anaerobic conditions for 6 hours. Meanwhile, the challenge plate consisting of the two gel treatments and MMBC-3 was prepared in a new microtiter plate in aerobic condition by adding 150 μl of each of the treatments and the medium in individual wells of the 96-well microtiter plate with the aid of 1 ml syringes. The 6-hour three-species biofilms or adherent cells present on the surface of the pegs were subjected to either MMBC-3 or Hyalugel^®^-ADO or blue^®^m oral gel for 1 hour in anaerobic condition. After the 1-hour treatment, the pegs were carefully lifted from the challenge plate and further incubated in the microtiter wells containing 200 μl of MMBC-3 for 24 hours in anaerobic conditions. After the 24-hour incubation, the 96-well microtiter plate comprised of bacteria that detached (as a consequence of either the treatment or biofilm formation on the pegs) and proliferated in planktonic form in the well. The pegs consisted of biofilm formed either due to the bacteria remaining on its surface post-treatment or the biofilm build-up due to the planktonic bacteria in the wells of the microtiter plate. Both samples (detached/planktonic cells and biofilm) are representative of the effectiveness of the treatment.

**Figure 1:**
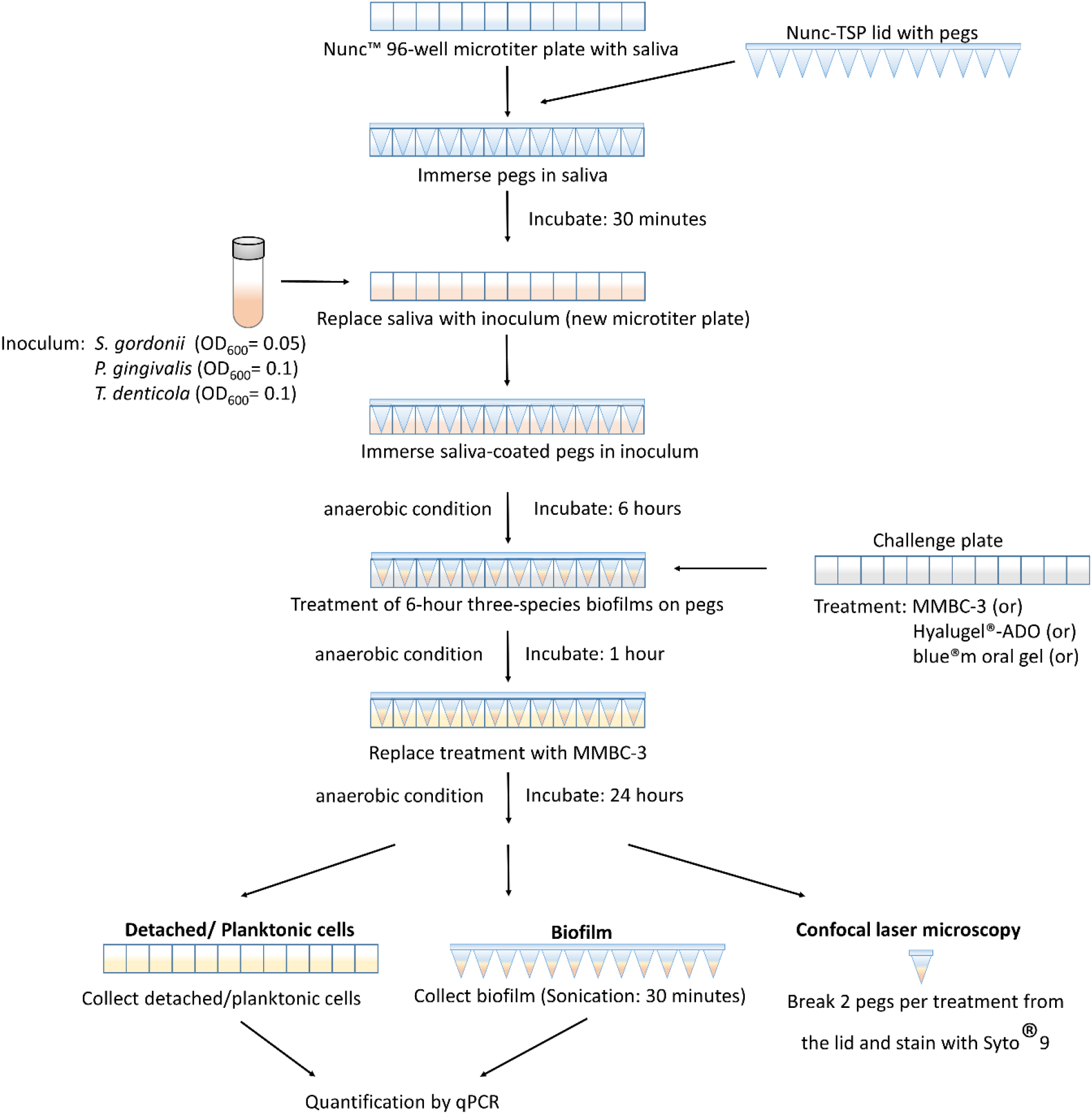
The process of testing antimicrobial gels on oral biofilms established on the surface of the pegs of the Calgary Biofilm Device.(see methods)

The detached/planktonic cells were collected from the microtiter plate and the biofilm were collected from the surface of the pegs by sonication (30 min) and were quantified by qPCR. 6 pegs (2 per condition) were broken with the help of pliers and stained with Syto^®^9 for visualization using confocal laser microscopy.

### Confocal microscopy and imaging

Treated and untreated biofilm-containing pegs after a 24-hour incubation in MMBC-3 were subjected to microscopic imaging after staining with 5 μM of Syto^®^9 green-fluorescent nucleic acid stain (Invitrogen, ThermoFisher Scientific) diluted in PBS. Briefly, 6 pegs were broken anaerobically from the lid with the help of sterile pliers and were placed in a microtiter plate containing the stain (200 μl) for 20 min. They were further transferred to Syto^®^9 stain-filled (200 μl) μ-slides (8 chambered coverslip, ibiTreat, Ibidi) anaerobically. The biofilm on the surface of the stained pegs was then observed *in situ* with a Leica TCS-SP5 confocal laser scanning microscope (Leica Microsystems, Wetzlar, Germany). An HC PL Apo 10X, 0.4 NA oil immersion objective lens was used for image capture and a numerical zoom of 2 was applied. The 488-nm UV diode and a 485 to 500-nm band-pass emission filter were used to detect all bacteria stained with Syto^®^9. Biofilm stacks (123 × 123 μm) acquired at 1 μm intervals were scanned with a line average of 2. Also, zoomed images were captured using HC PL Apo 63X, 1.4 NA oil immersion objective lens with a numerical zoom of 5.05.

Leica software (LAS AF V.2.2.1) was used for microscope piloting and image acquisition. Analysis of images based on Syto^®^9 was performed in ImageJ software V1.43m (National Institute of Health, USA) to obtain the maximum z-projection of the images.

### Quantification of bacteria by qPCR

The bacteria, consisting of *S. gordonii*, *P. gingivalis* and *T. denticola*, in the initial inoculum and the planktonic/detached cells collected after treatment were centrifuged (8000xg, 20°C, 10 min) and the pellets were resuspended in 150 μl of Lysis buffer (20 mg/ml lysozyme in 20 mM Tris-HCl, 2 mM EDTA, 1.2 % Triton X, PBS, pH 8). Biofilms were collected in 150 μl Lysis buffer by sonication for 30 min using a water bath sonicator (Ultrasonic cleaner). The biofilm collected from 3 pegs from the same condition (treated or untreated) were pooled together to increase the sample volume and to compensate for the low biofilm surface area on the pegs (approximately 44 mm^2^ per peg) (Harrison *et al.*, 2010; MBEC assay procedural manual, version 2.0, Innovotech). All samples were subjected to DNA extraction using QIAamp DNA Mini kit (Qiagen) according to the manufacturer’s instructions with slight modification i.e. the lysis using proteinase K was performed overnight.

Quantitative PCR was performed in a total reaction volume of 12.5 μl containing 6 μl Takyon™ Low Rox SYBR^®^ MasterMix dTTP Blue (Eurogentec), 0.5 μl of each primer (5μM), and 1 μl of the sample. Amplification of the extracted DNA template was performed in QuantStudio™ 7 Flex Real-Time PCR System (Applied Biosystems) by initial incubation of 2 min at 55°C and 10 min at 95°C, followed by 40 cycles of 15 sec at 95°C and 1 min at 60°C. A melt curve stage was performed consisting of 15 sec at 95°C followed by a temperature gradient from 60°C to 95° C with fluorescence measured in an increment of 1°C every 15 sec.

The concentrations of the DNA samples were determined in comparison with the defined concentrations of DNA standards set in the range of 0.0001 to 10 ng with purified genomic DNA from each of the three species. Primers used were specific to each species targeting the 16S ribosomal RNA taking into account specific genome weights (2.58 × 10^−6^ ng for *P. gingivalis*, 3.12 × 10^−6^ ng for *T. denticola* and 2.41 × 10^−6^ ng for *S. gordonii*) (Ammann *et al.*, 2013; Martin *et al.*, 2017). The primers used in this study are listed in Table 1.

**Table 1:**
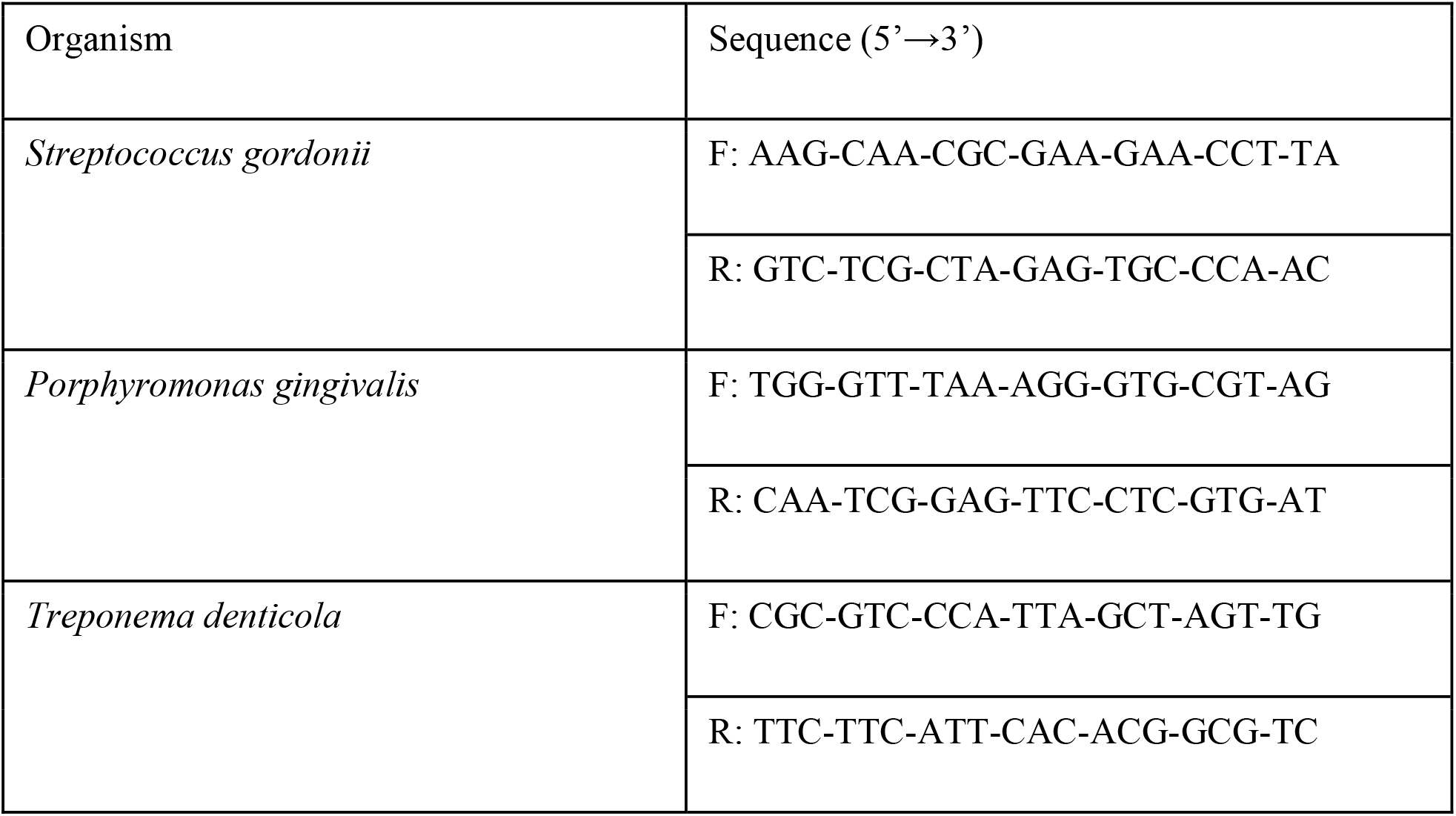
Species-specific primer sequences used in this study.

### Statistical analysis

All the experiments were done with 3 biological and 3 technical replicates. Statistical analysis was performed using the two-tailed unpaired student t-test and a p-value of less than 0.05 was considered statistically significant.

## RESULTS

### Establishment of the three-species oral biofilm on the peg-lid using MMBC-3 medium

The method of using Nunc-TSP lids with pegs allowed the formation and growth of the three-species oral biofilm on the surface of the pegs. Using this method, biofilm that formed on the peg surface after 31 hours (6 hours of growth prior to treatment + 1 hour of treatment + 24 hours of post-treatment growth, see methods and Figure 1) in MMBC-3 medium were quantified by qPCR (Figure 3A) and visualized by confocal fluorescent microscopy (Figure 2A and D). The microscopic images are representative of the total biofilm density and do not differentiate individual species. However, it enables the visualization of clusters of bacteria (Figure 2D) when zoomed. Additionally, the planktonic or detached bacteria in the 96-well microtiter plate were also quantified by qPCR (Figure 3A). The concentration of each species (*S. gordonii*, *P. gingivalis* and *T. denticola*) in the biofilm (Figure 3B) and planktonic/ detached condition (Figure 3C) were also measured.

**Figure 2:**
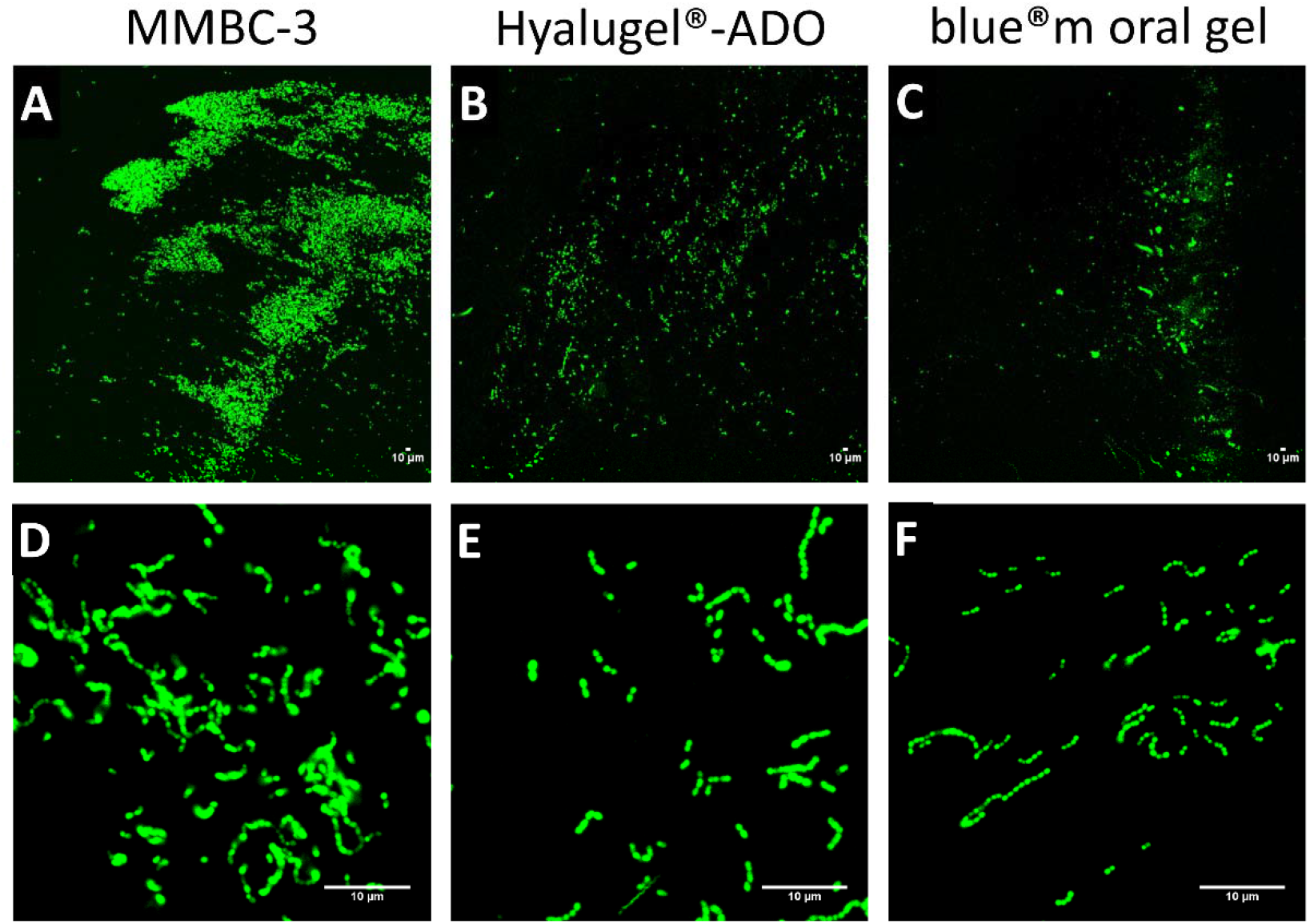
Representative microscopic images of the three-species oral biofilm on the surface of the pegs. Post 1-hour treatment (with MMBC-3, Hyalugel^®^-ADO or blue^®^m oral gel) of the 6-hour biofilms on the surface of the pegs, the pegs were incubated in MMBC-3 for 24 hours. The pegs were further broken from the lid using pliers and stained using Styo^®^9. The stained pegs were visualized using the Leica TCS-SP5 confocal laser scanning microscope. To compare between the three treatments, maximum z-production of the Z stack were taken using 10X oil immersion objective lens and numerical zoom of 2: (A), (B), (C). Magnified images were captured using the 63X oil immersion objective lens, numerical zoom of 5.05: (D), (E), (F).

**Figure 3:**
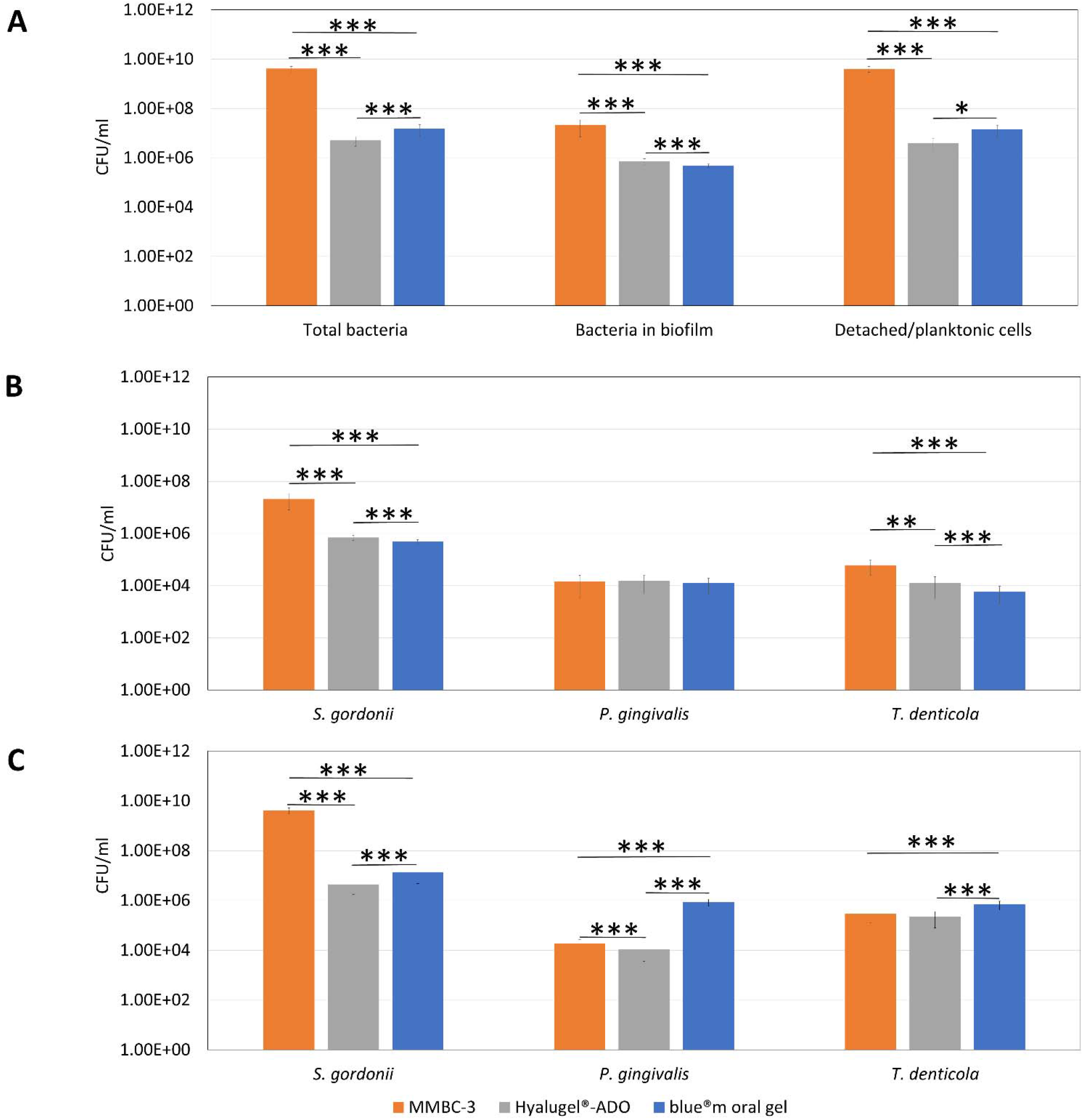
qPCR quantification of the number of bacteria (CFU/ml) in the biofilm (collected from the pegs) and in planktonic form (collected from the 96-well microtiter plate). The 6-hour three-species biofilm on the pegs were subjected to MMBC-3 or Hyalugel^®^-ADO or blue^®^m oral gel for 1 hour and further incubated in MMBC-3 for 24 hours. The planktonic cells were collected from the 96-well plate while the biofilm was collected from the pegs and quantified. (A) Total concentration (CFU/ml) of bacteria (planktonic/detached + biofilm), bacteria in biofilm and bacteria detached or in planktonic form after each treatment. (B) Concentration of each species in the biofilm after each treatment. (C) Concentration of each species in planktonic/ detached cells for each treatment. p-value < 0.05 = *, p-value < 0.01 = **, p-value < 0.001 = ***

The three-species biofilm quantified on the surface of the untreated pegs (3 pegs pooled together) is 2.08 × 10^7^ CFU/ml (Figure 3A). This value corresponds to 1.5 × 10^5^ CFU/mm^2^ (as the growth area per peg is 44 mm^2^). Martin *et al.*, (2018), in another study using the same species and medium reported biofilm formation of approximately 1 × 10^10^ CFU/ml in conventional μ-slides. This value corresponds to 1 × 10^8^ CFU/mm^2^ (as the growth area of each well in a μ-slides is 100 mm^2^). The concentration of the biofilm (in CFU/mm^2^) on the surface of the pegs is 650 times lesser than the concentration of biofilm on μ-slides. This difference is as expected since the pegs and μ-slides vary in structure and surface area. Also, conventional methods (like μ-slides) for biofilm formation pose the concerns of aggregation linked to sedimentation of bacteria in the wells. However, this concern can be disregarded in the case of peg-lids due to its protruding topology. Even though the concentration of bacteria in the biofilm on the surface of the pegs is less in comparison to conventional methods, it is still detectable by qPCR (in the species level) (Figure 3B) and can also be visualized by confocal microscopy (Figure 2A and D).

At the species level, the biofilm on the peg surface contains 99.2 % *S. gordonii*, 0.5 % *P. gingivalis* and 0.4 % *T. denticola* (Figure 4A). The composition of each species in the biofilm in conventional μ-slides (Martin *et al.*, 2018) is approximately 98.1 % *S. gordonii*, 1.3 % *P. gingivalis* and 0.6 % *T. denticola*. This ensures that even though the concentration of the biofilm is far less on the peg surface in comparison to μ-slides, it does not have a major effect on the overall composition (or ratio) of individual species for identical inocula. *S. gordonii* is seen to always predominate the population in the case of both peg-lids and μ-slides. This predominance of *S. gordonii* is also observed in the case of planktonic growth (Figure 4B). Further, on considering the sum of the bacteria in the biofilm and the detached (or planktonic) form as the total bacteria, we observe that a majority of bacteria remain in the planktonic form for all three species (Table 2).

**Table 2:** Species-wise percentage of biofilm and detached (or planktonic) cells post-treatment with either Hyalugel^®^-ADO or blue^®^m oral gel in comparison to MMBC-3.

|  | MMBC-3 |  | Hyalugel<sup>®</sup>-ADO |  | blue<sup>®</sup>m oral gel |  |
| --- | --- | --- | --- | --- | --- | --- |
|  | % biofilm | % detached/<br>planktonic<br>cells | % biofilm | % detached/<br>planktonic<br>cells | % biofilm | % detached/<br>planktonic<br>cells |
| *S. gordonii* | 0.4 ± 0.2 | 99.6 ± 0.2 | 15.2 ± 9.1 | 84.8 ± 9.1 | 3.5 ± 1.9 | 96.5 ± 1.9 |
| *P. gingivalis* | 35.0 ± 13.9 | 65.0 ± 13.9 | 51.6 ± 22.8 | 48.4 ± 22.8 | 1.2 ± 1.1 | 98.8 ± 1.1 |
| *T. denticola* | 12.8 ± 3.6 | 87.2 ± 3.6 | 5.7 ± 4.1 | 94.3 ± 4.1 | 0.7 ± 0.4 | 99.3 ± 0.4 |

100% stands for the sum of the percentages of bacteria in the biofilm and the detached (or planktonic) form.
% Calculation:
Percentages of bacteria in the biofilm:
Percentages of bacteria in the detached (or planktonic) form:

**Figure 4:**
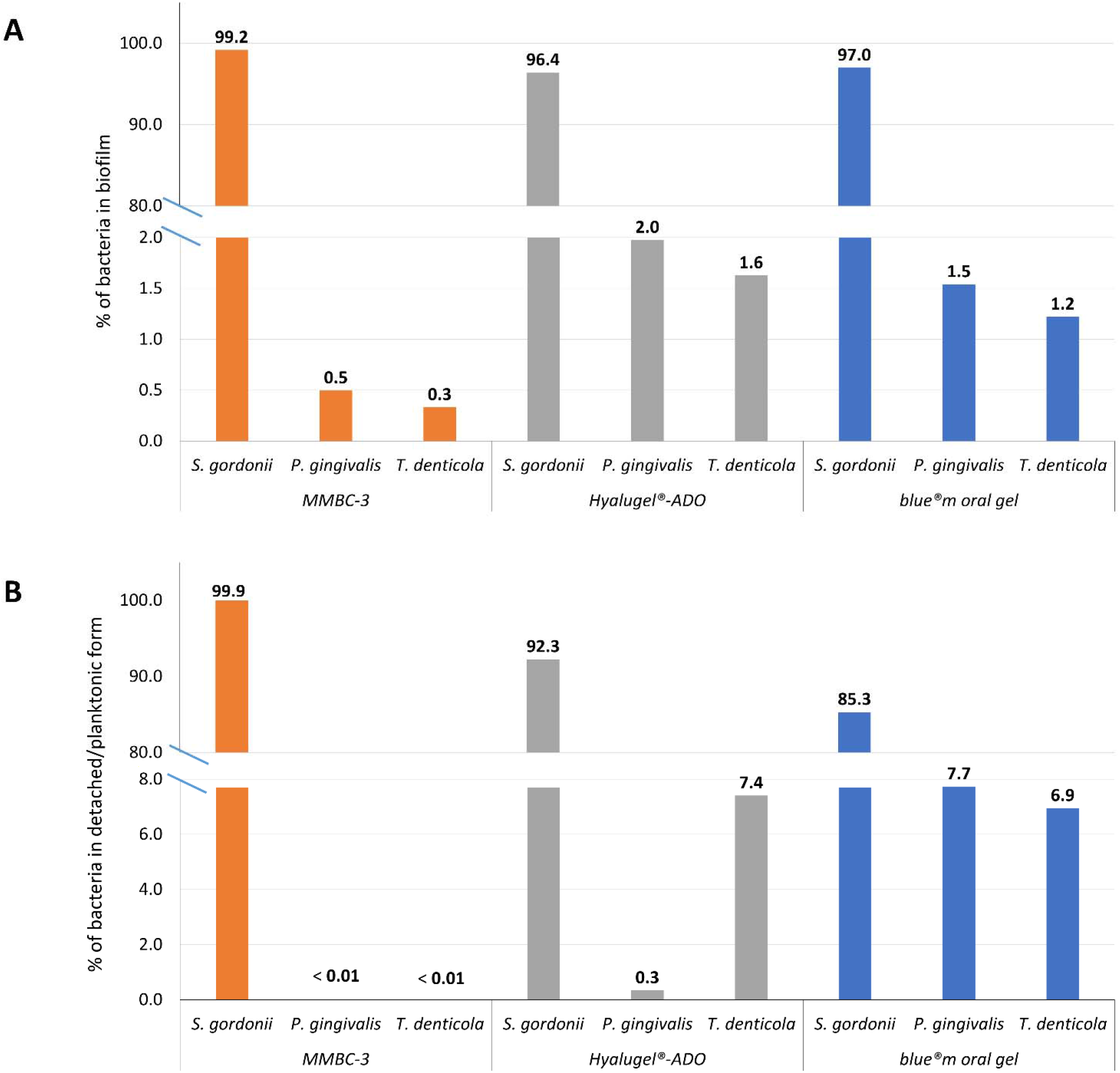
Composition of each species in the biofilms and in the detached/planktonic form post-treatment. Percentage of each species (*S. gordonii*, *P. gingivalis*, *T. denticola*) in the biofilms (A) and in the detached/planktonic form (B) after treatment with either Hyalugel^®^-ADO (grey bars) or blue^®^m oral gel (blue bars) in comparison to MMBC-3 (orange bars). The percentage of each species is mentioned above respective bars. 100% stands for the sum of the percentages of the three species either in the biofilm or the detached (or planktonic) form.

This method thus enabled the establishment of an oral biofilm model consisting of a facultative anaerobic commensal and two anaerobic pathogens on the surface of the pegs. It further allowed the analysis of the composition and quantification of each species in the biofilm and planktonic form. The use of the parameters described above will permit the comparison between various antibacterial gels against an oral biofilm model.

### Effect of antibacterial gels against a 6-hour three-species oral biofilm model

Next, in order to validate this method (Figure 1) on gel formulations, we tested the effect of two commercially available gels, namely, Hyalugel^®^-ADO and blue^®^m oral gel on the 3-species 6-hour biofilm established using MMBC-3 on the surface of the pegs. The two gels have comparable viscosities, 35000-60000 cP for Hyalugel^®^-ADO, and 25000-50000 cP for blue^®^m oral gel, as given by respective manufacturers. No gel without potential active compound was available to discriminate between the effect of viscosity and the effect of the active compound. Therefore, the method used evaluated both parameters against oral biofilm and enabled the comparison of gels with comparable viscosity.

The method allowed us to assess the number of sessile and planktonic bacteria remaining after treatment. We could also identify the most impacted species amongst a mixed biofilm. Focusing on the two gels used to implement the method, the concentration of the total bacteria was significantly reduced from 4.11 × 10^9^ CFU/ml to 5.15 × 10^6^ CFU/ml after treatment of the 6-hour three-species biofilm with Hyalugel^®^-ADO in comparison to MMBC-3 (Figure 3A). The decrease is evident in the concentration of bacteria in the biofilm as well as the concentration of bacteria in the detached/planktonic form. A similar trend was observed in the case of blue^®^m oral gel with total number of bacteria decreasing from 4.11 × 10^9^ CFU/ml to 1.50 × 10^7^ CFU/ml when compared to MMBC-3. The treatment with blue^®^m oral gel showed a decrease in the concentration of bacteria in the biofilm but an increase in detached/planktonic cells when compared to the treatment with Hyalugel^®^-ADO (Figure 3A). The blue^®^m oral gel showed greater efficiency in decreasing the concentration of bacteria in the biofilm when compared to Hyalugel^®^-ADO (Figure 3A). This decrease in biofilm concentration from MMBC-3 to Hyalugel^®^-ADO to blue^®^m oral gel is also evident in the microscopic images (Figure 2A, B, C).

The method used was efficient to monitor the variations in the concentration of each species after treatment. In the present protocol used to implement the method, the concentration of bacteria in the biofilm decreased significantly in the case of *S. gordonii* and *T. denticola* from MMBC-3 to Hyalugel^®^-ADO to blue^®^m oral gel while no effect of the two gels was seen on *P. gingivalis* (Figure 3B). Alternately, the concentration of *S. gordonii* and *P. gingivalis* in the detached/planktonic form, post-treatment with Hyalugel^®^-ADO decreased significantly in comparison to MMBC-3 (Figure 3C). No effect of Hyalugel^®^-ADO in comparison to MMBC-3 was observed on the concentration of *T. denticola* in the detached/planktonic form. In contrast, post-treatment with blue^®^m oral gel, the concentration of detached/planktonic *P. gingivalis* and *T. denticola* significantly increased in comparison to MMBC-3, while that of *S. gordonii* decreased. Also, the treatment with blue^®^m oral gel showed a higher concentration of all three species in the detached/planktonic form when compared to treatment with Hyalugel^®^-ADO.

Therefore, this method permitted the analysis of the effect of treatment on the composition of the biofilm and planktonic cultures. The percentage of *S. gordonii* in the antibacterial gel-treated (Hyalugel^®^-ADO and blue^®^m oral gel) biofilms decreased in comparison to untreated biofilm (MMBC-3) while the percentage of *P. gingivalis* and *T. denticola* increased post-treatment (Figure 4A). However, the ratio of the three species in the biofilm remained similar between the two treatments. The percentage of detached/planktonic *S. gordonii* decreased from untreated to treated biofilms, the least percentage being in the case of blue^®^m oral gel (Figure 4B). As a result, the percentage of the planktonic form of *P. gingivalis* and *T. denticola* increased post-treatment when compared to untreated biofilms. The percentage of planktonic *P. gingivalis* is higher in the case of blue^®^m oral gel-treated biofilms while the percentage of planktonic *T. denticola* is similar between the two treatments.

Finally, using this method, it is possible to evaluate the effect of treatment on the distribution between sessile and planktonic cells for each species. The species-wise percentage of biofilm and detached planktonic cells post-treatment with the gels in comparison to MMBC-3 was evaluated, where 100% constituted the sum of the percentages of bacteria in the biofilm and the detached planktonic form. The percentage of biofilm post-treatment with Hyalugel^®^-ADO in comparison to MMBC-3, was higher in the case of *S. gordonii* and *P. gingivalis* while lower in the case of *T. denticola*. Treatment with blue^®^m oral gel only modified and increased the percentage of *S. gordonii* in the biofilm in comparison to untreated biofilms. The percentage of planktonic bacteria was significantly reduced in the case of *S. gordonii* between untreated and treated biofilms (from 99.6 % for MMBC-3 to 96.4 % for blue^®^m oral gel to 84.8 % for Hyalugel^®^-ADO). For planktonic form of *P. gingivalis* a reduction was observed when Hyalugel^®^-ADO-treated biofilms were compared to MMBC-3 (from 65.0 % for MMBC-3 to 48.4 % for Hyalugel^®^-ADO). However, an increase in the percentage of planktonic form of *P. gingivalis* was observed in the case of blue^®^m oral gel-treatment as compared to Hyalugel^®^-ADO and MMBC-3. Also, an increase in the percentage of planktonic form of *T. denticola* was seen after gel treatment (from 87.2 % for MMBC-3 to 94.3 % for to Hyalugel^®^-ADO to 99.3 % for blue^®^m oral gel).

In short, the method was therefore efficient to determine the specific effect of both treatments on the oral biofilm: Hyalugel^®^-ADO (containing 0.2 % hyaluronic acid), reduced the planktonic growth of *S. gordonii* and *P. gingivalis* while it did not affect planktonic growth of *T. denticola*. This is in agreement with previously reported results (Ardizzoni *et al.*, 2011; Pirnazar *et al.*, 1999). Further, Hyalugel^®^-ADO reduced the biofilm formation of *S. gordonii* and *T. denticola* but did not affect the biofilm growth of *P. gingivalis* when compared to untreated biofilms. blue^®^m oral gel in comparison to untreated biofilms, reduced the overall growth of *S. gordonii*, increased the detached planktonic growth of *P. gingivalis* and *T. denticola* and reduced the biofilm growth of *S. gordonii* and *T. denticola*.

## DISCUSSION

Traditional methods for the development of biofilms (like microtiter plates, μ-slides, Ludin chambers etc.) can pose difficulties in testing the antimicrobial effect of gels against biofilms. In this study, we developed a high throughput method to test antibacterial gels against multispecies oral biofilm. This method uses the principle of the Calgary Biofilm Device to grow biofilms and was adapted for the growth of multispecies biofilm consisting of oral bacteria. We used the MMBC-3 (Martin *et al.*, 2018) as the growth medium especially designed for the growth of the three species used in this study. We use an apparatus/arrangement consisting of a 96-well microtiter plate and a lid with pegs. With the help of the microtiter plate, biofilms are established on the surface of the pegs. These biofilms are further challenged with antibacterial gels to test the effect of these gels against the oral biofilm model. Our method resolves the concerns posed by conventional methods by growing the biofilm on pegs while preparing the treatment in a microtiter plate. The separation of the treatment from the biofilm is thus not needed. We have devised a means of analyzing the effect of the treatment by further incubating the treated biofilm in a fresh microtiter plate containing medium (MMBC-3). Here, we assess the ability of the bacteria in the treated biofilm to further grow as biofilm (on the peg-surface) or planktonic culture (in the microtiter plate) in the MMBC-3 medium. Briefly, our method nullifies the need for separation of bacteria (either in biofilm or planktonic form) from the treatment/gel while it also allows the analysis of each species in the biofilm and planktonic form post-treatment. It is known that gels have an inherent shear force due to their viscous and adhesive nature which can result in the removal of bacteria from the surface of the pegs. In this study, to test the method, we compare the efficiency of two commercially available gels (with comparable viscosity) in biofilm reduction as a combined effect of its inherent shear force and its respective active compounds and against untreated biofilms.

The reduction in the biofilm in comparison to untreated biofilms may be due to the combined effect of the active antimicrobial compound (hyaluronic acid, sodium perborate or lactoferrin) and the viscous nature of gels. It cannot be excluded that since the 6-hour biofilm is scanty, its removal from the peg surface is enabled by the inherent viscous/adhesive nature of the gel. The method allowed us to compare two treatments with comparable viscosity. Here, blue^®^m oral gel induced higher concentration of the three species as planktonic cells while lower *S. gordonii* and *T. denticola* concentrations in the biofilm, compared to Hyalugel^®^-ADO. The difference may be due to active compounds as both gels have comparable viscosity. The oxygen donor present in blue^®^m oral gel may have an effect on the obligate anaerobe *T. denticola* or boost the peroxygenic activity of *S. gordonii* and change the biofilm behaviour (Chathoth *et al.*, 2020). Lactoferrin of blue^®^m oral gel can reduce the initial attachment of *S. gordonii* (Arslan *et al.*, 2009). This may explain the decrease in *S. gordonii* in blue^®^m oral gel-treated biofilms.

Our method demonstrated the ability of both gels in the reduction of the overall biofilm growth in comparison to untreated biofilm. They were especially effective in reducing *S. gordonii* and *T. denticola* in the biofilm. *S. gordonii* along with 16 other genera, has been previously classified under ‘signatures of dysbiosis’ due to its predominance in patients with periodontitis and edentulism (Hunter *et al.*, 2016). Also, its role in co-aggregation and metabolic interactions with other periodontal pathogens is well-known (Hajishengallis and Lamont, 2016; Sakanaka *et al.*, 2015). On the other hand, *T. denticola*, a member of the red complex, is known for its metabolic symbiosis, co-aggregation and synergy with the keystone pathogen *P. gingivalis* (Ito *et al.*, 2010; Meuric *et al.*, 2013; Ng *et al.*, 2019). Hence, the reduction of *S. gordonii* and *T. denticola* in the oral biofilm model with a single treatment for 1 hour with either of the two gels is indeed a promising result. Both the gels have the potential of preventing oral dysbiosis and co-aggregation of periodontal pathogens. Hence multiple exposures/treatments per day with either of the two gels is likely to show better efficacy in the reduction of the dental plaque.

## CONCLUSION

Antimicrobial gels are a promising treatment and can additionally be used as dressings and fillers, particularly in periodontal pockets, where the pathogenic oral biofilm resides. This study describes the method of using peg-lids for testing antimicrobial gels on multispecies biofilms. It offers the possibility of simultaneous testing of multiple conditions with reproducible cell density (Goeres *et al.*, 2005). It eliminates concerns due to sedimentation of bacteria (which is possible in the case of conventional methods). The method presented in this study has been optimized for the growth and development of oral biofilms in an iron-controlled medium. The effect of gels against the oral biofilms is measured by analyzing the concentration of bacteria in the biofilm or planktonic form across treatments in comparison to the untreated biofilms. Another parameter that is assessed is the ratio of each species in the biofilm and planktonic form. Further, the ratio of the bacteria in the biofilm to the planktonic form is also evaluated to understand the effect of the treatment in comparison to untreated condition. Finally, the treated biofilms after incubation in MMBC-3 for 24 hours is also subjected to confocal laser microscopy to visualize the effect of the treatments on the biofilms. However, the number of the live and dead bacteria in the biofilm and planktonic growth after the treatments can be assessed to further optimize the method (Harrison *et al.*, 2007). Therefore, we will know if the antibacterial gels are bacteriostatic or bactericidal in action. Assays with various times of incubation of the biofilms prior to treatment can also be performed to model different pathogenic states. This method can be further adapted for other studies like testing antibacterial compounds (other than gels) against biofilms. Besides, biofilms other than oral biofilms can also be grown and studied using this method.

## AUTHOR STATEMENT

The study was designed by KC, BM and CB; experiments were performed by KC; the data was analysed by KC, BM and CB; the manuscript was written and edited by KC, BM, MBM and CB.

## ACKNOWLEDGEMENTS

We thank blue^®^m Europe B.V., Netherlands for sending us their product-blue^®^m oral gel, for this study and for product-related information. We also thank the manufacturer of Hyalugel^®^-ADO, Ricerfarma (Milan, Italy) and the distributer, Laboratoire COOPER, Melun, France, for providing the product-related information. Our sincere gratitude to the Microscopy Rennes Imaging Center (Biosit) for their technical assistance for the microscopy experiments.

